# Eukaryotic translation initiation factor 5A and its hypusination play an important role in proliferation and differentiation of the human enteric protozoan parasite *Entamoeba histolytica*

**DOI:** 10.1101/2020.08.21.260711

**Authors:** Ghulam Jeelani, Tomoyoshi Nozaki

## Abstract

The eukaryotic translation initiation factor 5A (eIF5A) is highly conserved and essential in all eukaryotes. However, the specific roles of eIF5A in translation and in other biological processes remain elusive. In the present study, we described the role of eIF5A, its posttranslational modifications (PTM), and the biosynthetic pathway needed for the PTM in *Entamoeba histolytica*, the protozoan parasite responsible for amoebic dysentery and liver abscess in humans. *E. histolytica* encodes two isotypes of eIF5A and two isotypes of enzymes, deoxyhypusine synthase (DHS), responsible for their PTM. Both of the two eIF5A are functional, whereas only one DHS (EhDHS1), but not EhDHS2, is catalytically active. The DHS activity increased ∽2000 fold when EhDHS1 was coexpressed with EhDHS2 in *Escherichia coli*, suggesting that the formation of a heteromeric complex is needed for full enzymatic activity. Both *EhDHS1* and *2* genes were required for *in vitro* growth of *E. histolytica* trophozoites, indicated by small antisense RNA-mediated gene silencing. In trophozoites, only *eIF5A2*, but not *eIF5A1*, gene was actively transcribed. Gene silencing of eIF5A2 caused compensatory induction of expression of *eIF5A1* gene, suggesting interchangeable role of two eIF5A isotypes and also reinforcing the importance of eIF5As for parasite proliferation and survival. Furthermore, using a sibling species, *Entamoeba invadens*, we found that *eIF5A1* gene was upregulated during excystation, while *eIF5A2* was downregulated, suggesting that *eIF5A1* gene plays an important role during differentiation. Taken together, these results have underscored the essentiality of eIF5A and DHS, for proliferation and differentiation of this parasite, and suggest that the hypusination associated pathway represents a novel rational target for drug development against amebiasis.

**Author summary:** Eukaryotic initiation factor 5A is a ubiquitous protein that is essential for cell proliferation. We examined the maturation, regulation, and function of eIF5A in *E. histolytica*. We found by small antisense RNA-mediated gene silencing that EhDHS1/2 and EheIF5A2 are essential for growth of *E. histolytica* trophozoites. We further found that only one eIF5A, EheIF5A2, of two isotypes was constitutively expressed in the trophozoites stage and silencing of *EheIF5A2* gene caused overexpression of the other eIF5A isotype (EheIF5A1) to partially rescue the growth defect in this parasites. Furthermore, we found that transcription of *eIF5A1* gene was stage-specifically upregulated during excystation in *E. invadens*. Taken together, we have demonstrated for the first time that the two eIF5As play important and distinct roles in *Entamoeba* biology. This study has also provided an answer to a long standing conundrum on the biological importance of polyamines: spermidine is essential for eIF5A hypusination essential for protein translation in *Entamoeba*. Our work should also help our understanding of the physiological significance of eIF5A and its post-translational modifications in other pathogenic eukaryotes and potentially lead to formulation of control measures against parasitic diseases.

## Introduction

Polyamines are low-molecular-weight nitrogenous bases that are essential for the regulation of cell growth and development [1]. Due to their polycationic nature, the polyamines have the ability to interact electrostatically with the majority of polyanionic macromolecules in cells and thereby influence a variety of processes including cell differentiation and proliferation, embryonic development, and apoptosis [2]. As a consequence, increased concentrations of the polyamines and their biosynthetic enzymes are observed in highly proliferating cells such as cancerous cells and parasitic organisms [3]. One of the demonstrated universal roles of polyamines, particularly spermidine, in eukaryotic cells is the formation of hypusine on the eukaryotic initiation factor 5A (eIF5A) (Fig 1). Hypusine is an unusual amino acid [N (ε)- (4-amino-2-hydroxybutyl)-lysine], which is uniquely synthesized on eIF5A at a specific lysine residue from spermidine by two catalytic steps [4]. This post-translational modification (PTM) in eukaryotes is achieved by the sequential reactions catalyzed by two enzymes: deoxyhypusine synthase (DHS) and deoxyhypusine hydroxylase (DOHH) (Fig 1) [4]. Hypusination is the most specific PTM known to date [5] and it is essential for eIF5A activity [6]. eIF5A is involved in elongation [7], termination [8], and stimulation of peptide bond formation [9], and it facilitates protein synthesis by resolving polyproline-induced ribosomal stalling; thus, its role seems indispensable in synthesis of proline repeat-rich proteins [10].

**Fig 1.**
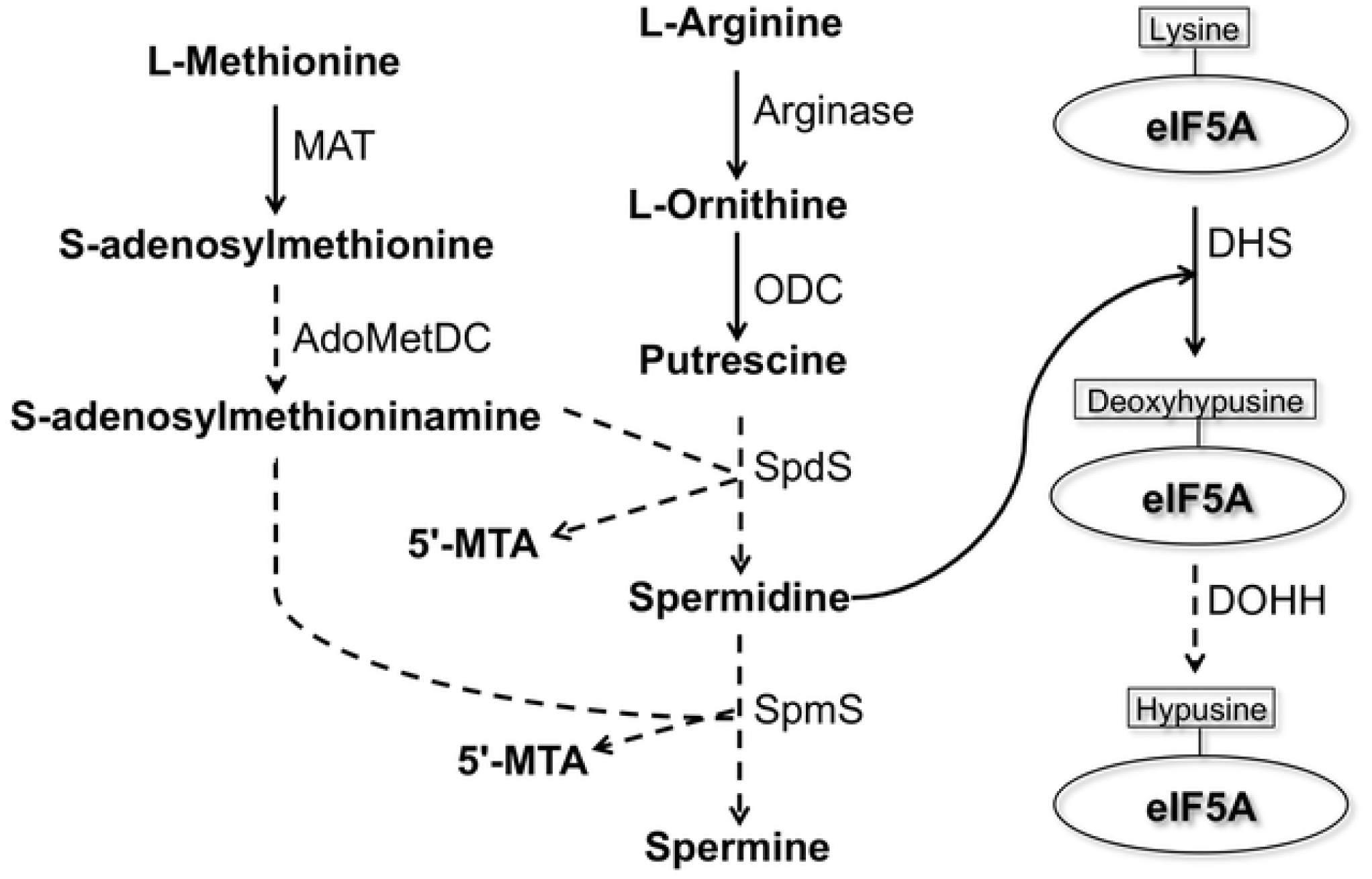
Presumed biosynthetic pathway of polyamines and deoxyhypusine/hypusine modifications on eukaryotic initiation factor 5A in *E. histolytica*. Solid lines represent the steps catalyzed by the enzymes whose encoding genes are present in the *E. histolytica* genome, whereas dashed lines indicate those absent or not yet identified. Abbreviations: 5’-MTA, 5’-methylthioadenosine; MAT, S-adenosylmethionine synthetase; ODC, ornithine decarboxylase, AdoMetDC, S-adenosylmethionine decarboxylase; SpdS, spermidine synthase; SpmS, spermine synthase; eIF5A, eukaryotic initiation factor 5A; DHS, deoxyhypusine synthase; DOHH, deoxyhypusine hydroxylase.

*E. histolytica* is a unicellular parasitic protozoan responsible for human amebiasis. The world health organization estimates that approximately 50 million people worldwide suffer from invasive amebic infections, resulting in 40 to 100 thousand deaths annually [11]. As a parasite, *E. histolytica* needs be able to cope with a wide variety of environmental stresses, such as fluctuation in glucose concentration, changes in pH, pO_2_, temperature, and host immune systems including oxidative and nitrosative species from neutrophils and macrophages during the life cycle [12]. Our previous metabolomic analyses have indicated that polyamines including putrescine, spermidine, and spermine are abundantly present in the proliferating and disease-causing trophozoites and the levels of these metabolites dramatically decrease during stage conversion from the trophozoites to the dormant cyst stage [13]. Interestingly, genome-wide survey of the reference genome (http://amoebadb.org/amoeba/) suggested that *E. histolytica* lacks a few key enzymes involved in polyamine biosynthesis, conserved in other bacterial and eukaryotic organisms: *S*-adenosylmethionine decarboxylase (AdoMetDC), spermine synthase (SpmS), and spermidine synthase (SpdS), suggesting that *E. histolytica* may possess a unique pathway or enzymes for polyamine biosynthesis, which can be further explored as a drug target against amebiasis.

Biosynthesis of spermidine and its requirement in the essential PTM of translation elongation factor eIF5A [14]. The *E. histolytica* genome revealed that this parasite possesses genes for deoxyhypusine synthase and eIF5A; however, *E. histolytica* seems to lack DOHH, which is involved in formation of mature hypusinated eIF5A (Fig 1). DHS has also been shown to be essential in all species where it was studied including mammals, yeast [15], and the kinetoplastids, *Trypanosoma brucei* [16] and *Leishmania* [17]. While most eukaryotes possess only a single *DHS* gene, *Entamoeba, Leishmania, and Trypanosoma* possess more than one *DHS* or *DHS-like* genes [18]. The presence of two *DHS* genes in these parasitic protozoa may be suggestive of unknown biological significance.

In the present study, we demonstrate that only one gene of the two *EhDHS* genes from *Entamoeba* encodes for the enzymatically active DHS. We show that the other DHS isotype encoded by the second gene is needed for a formation of a protein complex, required for maximal enzyme activity. We also show that both *EhDHS1* and *2* genes are required for optimal *in vitro* growth of *E. histolytica* cells. We have also found that only eIF5A2 protein, but not eIF5A1, is constitutively expressed in trophozoites and that silencing of *eIF5A2* gene causes compensatory expression of eIF5A1, suggesting that partially interchangeable roles of these proteins for growth or survival. Furthermore, we found that transcription of *eIF5A1* gene is upregulated during excystation while that of *eIF5A2* is downregulated, suggesting that eIF5A1 is involved in the differentiation. To date, this study represents the first case indicating that eIF5A and its hypusination plays an essential role in proliferation and differentiation of this pathogenic eukaryote.

## Results

### Identification and features of DHS and EIF5A genes and its encoded proteins from *E. histolytica*

While polyamines were previously demonstrated in trophozoites [13], their metabolic roles remained elusive. As hypusination utilizes spermidine as a substrate, we hypothesized that the amebic trophozoites utilizes spermidine for the PTM of eIF5A. A genome wide survey of eukaryotic DHS, which catalyzes the first step of hypusination of eIF5A, in the *E. histolytica* reference genome (AmoebaDB, http://amoebadb.org/amoeba/), by BLASTP analysis using human DHS as a query, revealed that *E. histolytica* possesses two possible DHS homologs, which showed 34% mutual identity. We designated EHI_098350 as EhDHS1 and EHI_006030 as EhDHS2. EhDHS1 contains the key catalytic lysine residue known to be conserved among orthologs from other eukaryotes, while the residue is not conserved in EhDHS2. Both EhDHS1 and EhDHS2 exhibits 47% amino acid sequence identity to human DHS (HsDHS), which is encoded by a single copy gene (Fig 2A). Multiple sequence alignment shows that EhDHS1 contains a complete set of the amino acid residues implicated for spermidine binding (4/4) that are conserved in HsDHS [19], whereas only two of four residues are conserved in EhDHS2. Nine and ten out of thirteen residues predicted to be involved in the NAD-binding are conserved in HsDHS are conserved in EhDHS1 and EhDHS2, respectively.

**Fig 2.**
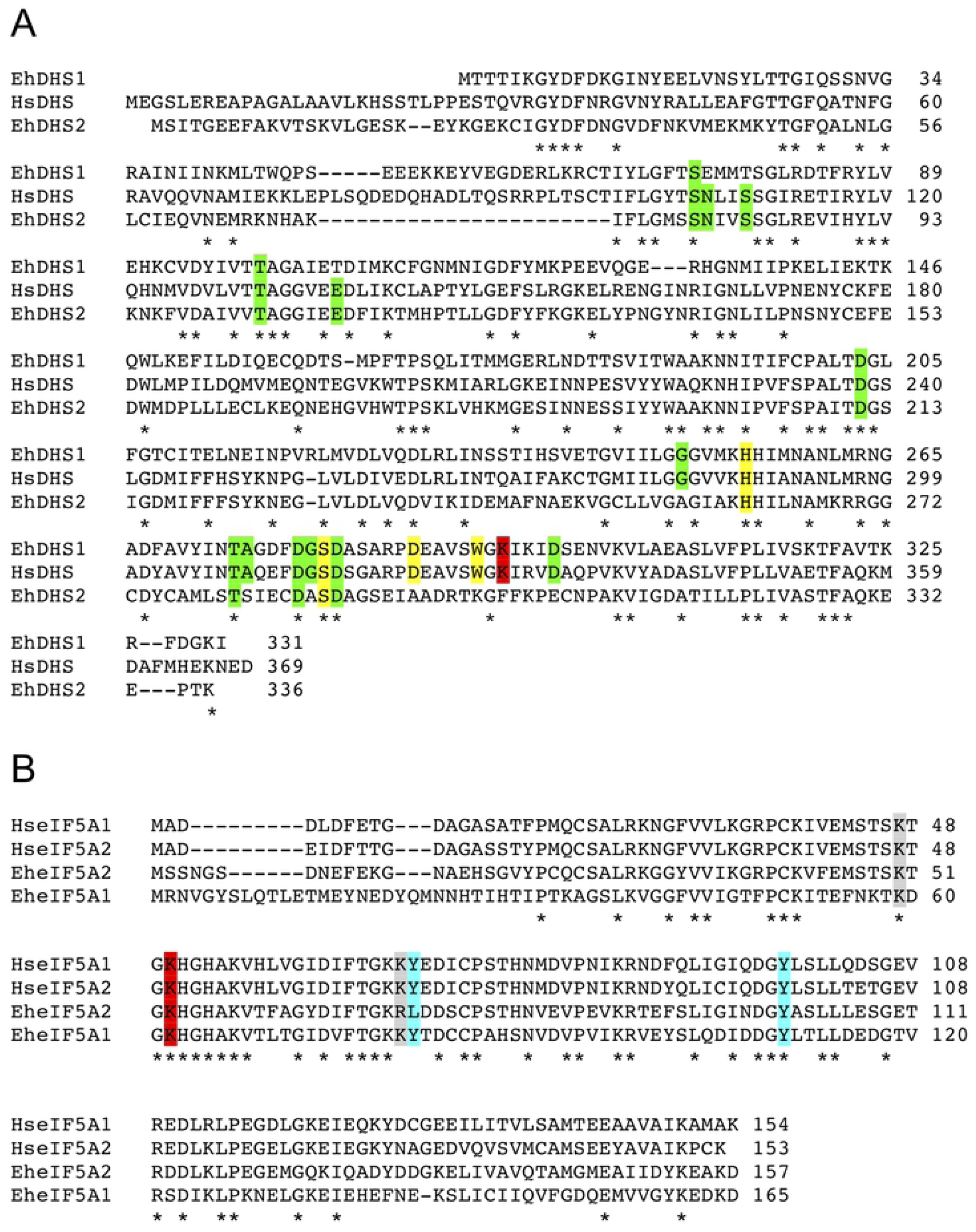
Multiple alignment of DHS and eIF5A protein sequences. The conserved residues are marked by asterisks (*). Sequence alignment was performed using ClustalW. **(A)** Alignment of DHS protein sequences from *E. histolytica* [EhDHS1 (XP_653614), EhDHS2 (XP_653426)], and *H. sapiens* (P49366). The catalytic lysine residue is shown on a red background, whereas NAD^+^ and spermidine binding sites are shown in green and yellow background, respectively. **(B)** Amino acid sequence alignment of *E. histolytica* and *H. sapiens* eIF5A isoforms. Accession numbers of these sequences are as follows: EheIF5A1 (XP_657374); EheIF5A2 (XP_651531); HseIF5A1 (P63241); and HseIF5A2 (AAG23176). The conserved lysines, which are expected to be hypusinated, are highlighted in red. The residues, which are post-transnationally modified by acetylation and sulfation in human eIF5A, are shown in grey and cyan background, respectively.

A genome wide survey identified four eIF5A proteins (EHI_151540, EHI_186480, EHI_177460, and EHI_151810). They are categorized into two groups (EheIF5A1 and EheIF5A2) based on mutual sequence similarity (S2 Table). We designated representative allelic isotypes as eIF5A1 for EHI_151540 (and EHI_177460) and eIF5A2 for EHI_186480 (and EHI_151810), respectively. A multiple alignment of two EheIF5A isotypes with two human eIF5A isoforms, by ClustalW program (http://clustalw.ddbj.nig.ac.jp/top-e.html), shows that the two glycine residues adjacent to lysine (red) are conserved in EheIF5A (amino acid positions 61 and 64 in eIF5A1; 52 and 55 in eIF5A2) (Fig 2B). This Gly-X-Y-Gly motif was reported to be critical for β turn structure and the proper orientation of the deoxyhypusine/hypusine side chain [20]. Lysine residues at positions 62/53 and 59/50 in eIF5A1/2, respectively, are presumed to be the sites for hypusination and acetylation, respectively [21]. In addition, tyrosine residues at positions 81 and 110 in eIF5A1 and at position 101 in eIF5A2 are likely the sites of sulfation [22]. The highly conserved hypusination domain in EheIF5A indicates an early establishment of the domain function and likely its PTMs throughout eukaryotic evolution.

### Recombinant EhDHS1 is active against eIF5A1/A2 while EhDHS2 is inactive

To understand the role of two isotypes of EhDHS, enzymological characterization of the recombinant EhDHS1 and 2 was performed. The recombinant EhDHS1, EhDHS2, EheIF5A1, and EheIF5A2 were produced at the level of 2.0-2.5% of the total soluble proteins in *E. coli* BL21 cells. SDS-PAGE analysis followed by Coomassie Brilliant Blue staining showed that the purified recombinant EhDHS1, EhDHS2, EheIF5A1, and EheIF5A2 proteins were present as apparently homogenous 37.2, 37.4, 18.6 and 17.1 kDa proteins, respectively, under reducing conditions (S1A Fig). The mobility of all recombinant proteins was consistent with the predicted size of the monomeric proteins with an extra 2.6 kDa histidine tag added at the amino terminus. We next tested the enzymatic activities of recombinant EhDHS1 and EhDHS2 using recombinant EheIF5A1 and EheIF5A2 as substrates. We found that EhDHS1 is active with the approximate specific activity of 13.0±1.0 pmol/min/mg with EheIF5A1 and 9.4±2.3 pmol/min/mg protein with EheIF5A2 (Fig 3B). The observed enzyme activity was 1000-2000 fold lower than that of DHSs from other organisms [19,23], which may be explained in part by the fact that key residues involved in substrate binding are missing in EhDHS1. In contrast, recombinant EhDHS2 showed no detectable activity (Fig 3B), which is consistent with the fact that EhDHS2 lacks key amino acid residues including lysine implicated for the formation of enzyme-substrate intermediates and the two amino acids involved in the spermidine binding.

**Fig 3.**
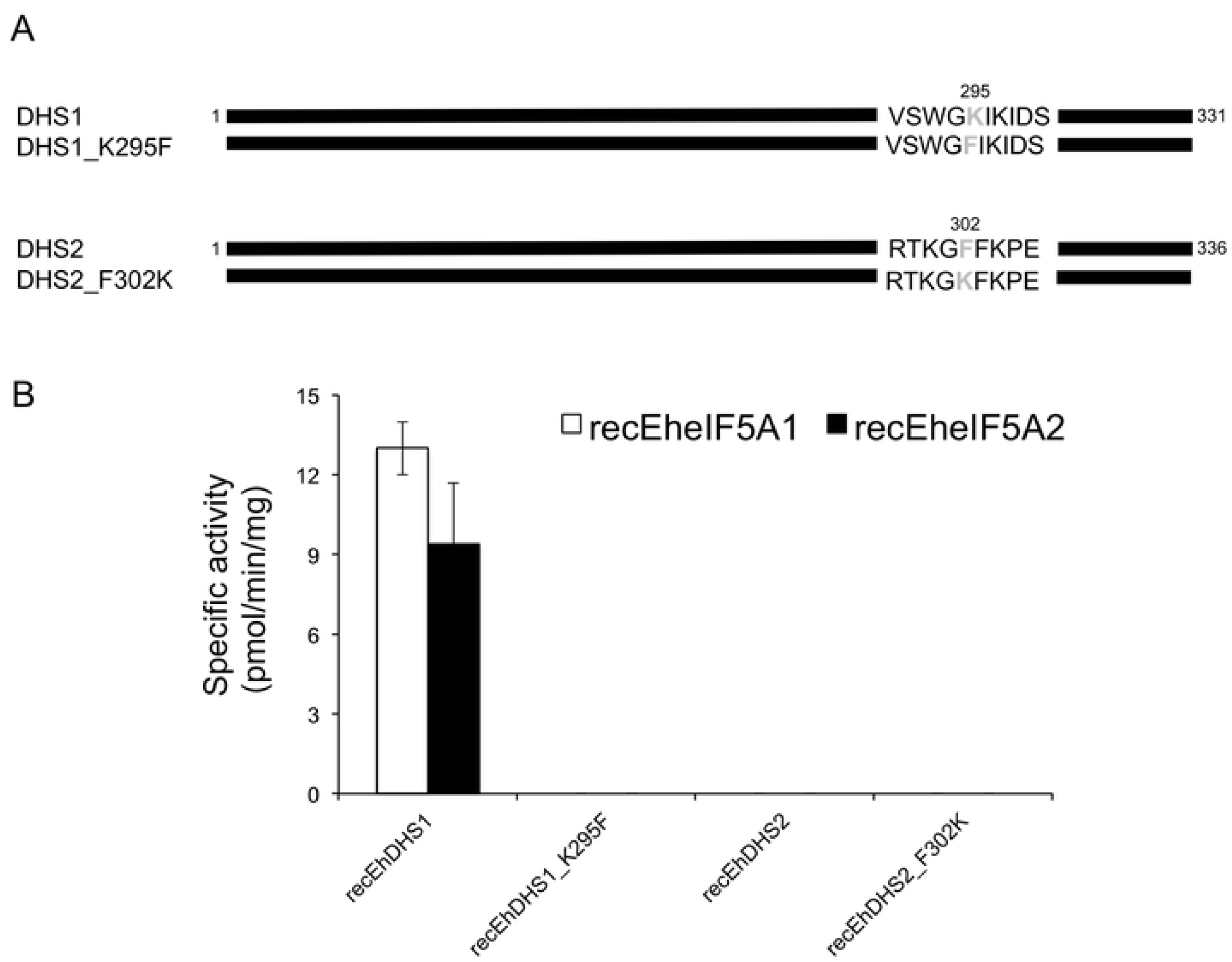
Schematic diagram of wild type and mutant forms of EhDHS1 and EhDHS2, and characterization of EhDHS1 and EhDHS2 mutants. **(A)** Schematic representation of DHS1 and DHS2 mutant. Amino acid residues mutated are shown in grey. Amino acid positions are also shown. **(B)** Enzymatic activity of recombinant EhDHS1, EhDHS2, EhDHS1_K295F, and EhDHS2_F302K using EheIF5As substrates. The mean ± S.D. of three independent experiments performed in triplicate is shown.

### The conserved lysine residue in the catalytic site is necessary, but not sufficient for DHS enzymatic activity

To better understand the physiological role of the apparently redundant, active and inactive EhDHS isotypes in *E. histolytica*, and, more specifically, to understand the reason for the lack of activity of EhDHS2, we made two EhDHS1 and EhDHS2 mutants, in which the lysine residue implicated for catalysis is replaced with phenylalanine in EhDHS1 or phenylalanine was replaced with lysine in EhDHS2, by site-directed mutagenesis (EhDHS1_K295F and EhDHS2_F295K) (Fig 3A, S1B Fig) Both the mutants were inactive (Fig 3B) with either EheIF5A1 or EheIF5A2 as a substrate, suggesting that the lysine residue in the enzyme-substrate intermediate site is essential for activity, but not sufficient, and that substitutions of other amino acids responsible for spermidine and NAD^+^ binding also contribute to the lack of enzymatic activity in EhDHS2.

### EhDHS1 forms a complex with EhDHS2, which strongly enhances enzymatic activity of EhDHS1

We speculated if enzymatically inactive EhDHS2 plays a regulatory role of the enzymatic activity harbored by EhDHS1, as previously reported *for* DHS from *Trypanosoma brucei* [16]. We used a system that allows coexpression of two proteins in *E. coli* as described previously [24]. Using pETDuet-1 vector, we were able to create a bacterial strain that expresses EhDHS1 with the amino-terminal His-tag and EhDHS2 with the carboxyl-terminal S-tag. Co-expression of EhDHS1 and EhDHS2 was verified by immunoblot analysis using anti-His and anti-S tag antibodies (Fig 4A). Recombinant His-tag DHS1 and S-tag DHS2 were copurified using Ni^2+^-NTA resins as described in material and methods section. The purified proteins were present in approximately equimolar amounts, indicating that EhDHS1 and EhDHS2 form a stable complex with a ratio of one to one. DHS activity of the dimeric EhDHS1/EhDHS2 proteins was measured using EheIF5A1 or EheIF5A2 as substrates. We found a more than 2,000 fold increase in the enzymatic activity when EhDHS1 and EhDHS2 were coexpressed, as compared to EhDHS1 alone (26.1±6.0 μmol/min/mg protein with EheIF5A1 and 22.4±10.0 μmol/min/mg protein with EheIF5A2) (Fig 4B).

**Fig 4.**
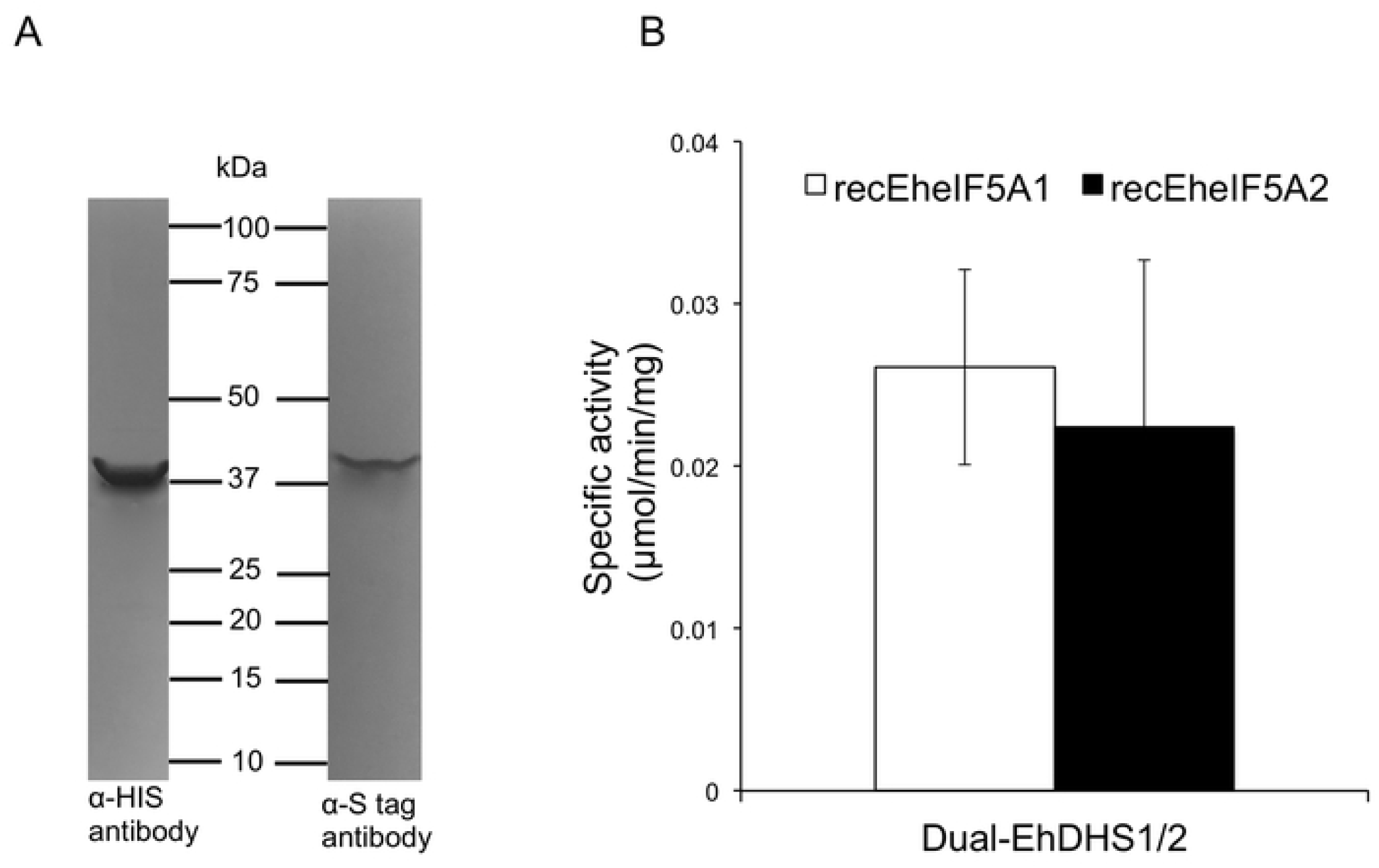
In vitro complex formation between EhDHS1 and EhDHS2 resulted increase in enzyme activity. **(A)** Immunoblot analysis of coexpression of EhDHS1, which contains the amino-terminal His-tag, and EhDHS2, which contains the carboxyl-terminal S-tag, using anti-His and anti-S tag antibodies. **(B)** Specific activity of coexpressed recombinant EhDHS1 and EhDHS2 using EheIF5A1 and EheIF5A2 as substrates. The means ± standard deviations of three independent experiments performed in triplicate are shown.

### Both *EhDHS1* and *EhDHS2* genes appear to be essential for *E. histolytica* growth

To investigate the role of DHS1 and DHS2 in *E. histolytica*, we exploited antisense small RNA-mediated epigenetic gene silencing to repress the *EhDHS1* and *EhDHS2* genes in *E. histolytica* G3 strain [25, 26]. However, our repeated attempts to create a transformant in which *EhDHS1* gene expression was repressed by gene silencing failed, suggesting the essentiality of the gene. In contrast, we successfully gene silenced *EhDHS2* (Fig 5A) and found that *EhDHS2* gene-silenced transformants displayed asevere growth defect (Fig 5B). Interestingly, we found that in *EhDHS2* gene silenced strain the steady-state levels of transcripts of *EhDHS1* and *EheIF5A1* were up regulated approximately 4 fold (Fig 5A). We next checked whether *DHS2* gene silencing affects the hypusination levels in the cells. Using anti-hypusine polyclonal antibody [27] we examined the hypusination levels in *EhDHS2* gene silenced and control transformants. We found that hypusination levels decreased by 64% (Fig 5C), suggesting that EhDHS2 plays, likely via interaction with and activation of EhDHS1, an important role in maintaining the hypusination levels in the cell.

**Fig 5.**
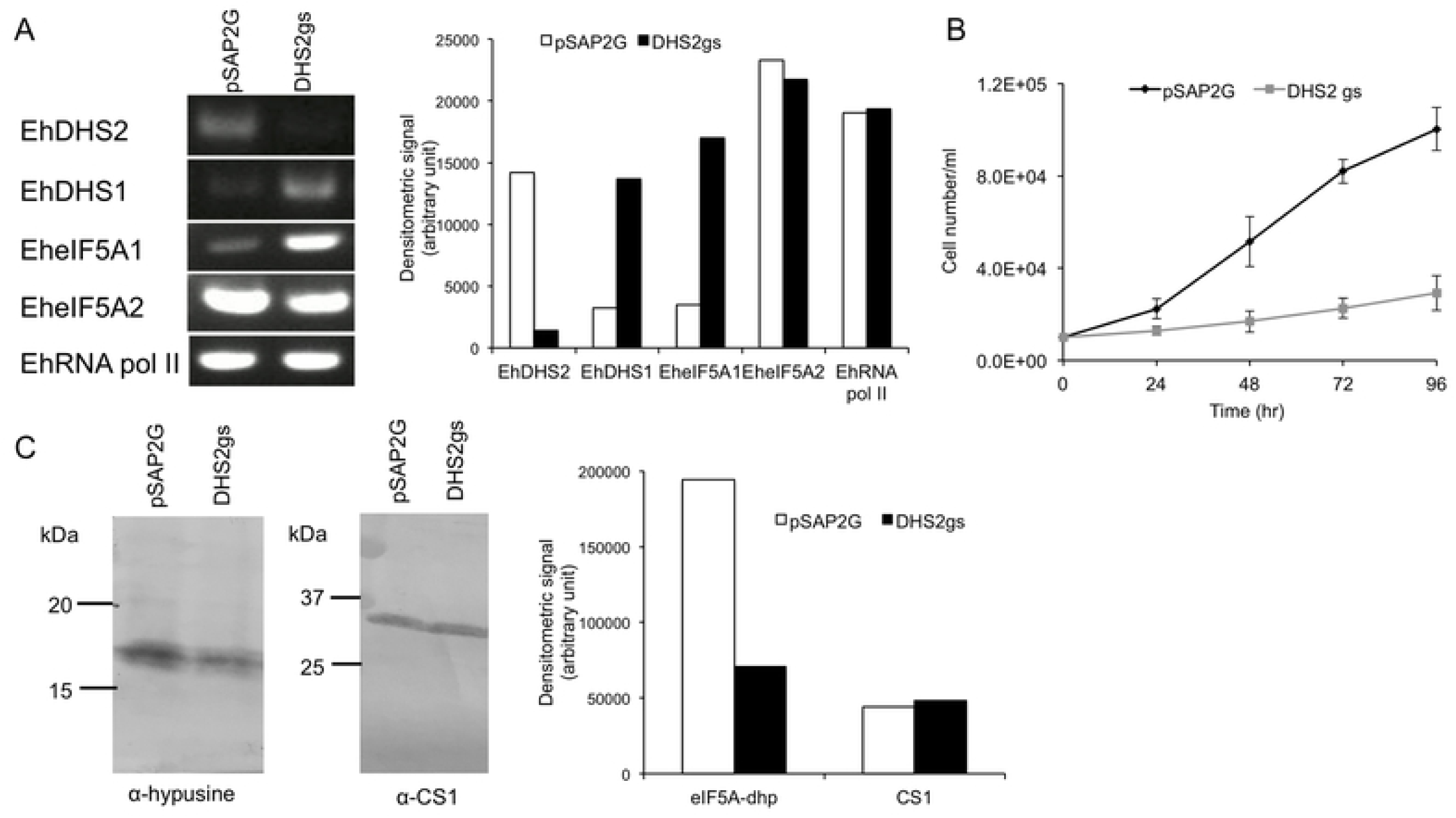
Effects of *EhDHS2* gene silencing on gene expression of related genes and growth of*E. histolytica* trophozoites. **(A)** Evaluation of gene expression by semi-quantitative RT-PCR analysis of *EhDHS2* gene silenced transformant. The steady-state levels of transcripts of *EhDHS1, EhDHS2, EheIF5A1, EheI5FA2* and *EhRNA pol II* genes were measured in trophozoites of G3 strain transfected with either empty vector (psAP2G) or the *EhDHS2* gene silencing plasmid (psAP2G-DHS2). cDNA from the generated cell lines (psAP2G and DHS2gs strains) was subjected to 30 cycles of PCR using specific primers for the *DHS2, DHS1, eIF5A1, eI5FA2* and *RNA pol II* genes. RNA polymerase II served as a control. PCR analysis of samples without reverse transcription was used to exclude the possibility of genomic DNA contamination. The densitometric quantification of the bands, shown in the right graph, was performed by Image J software, and the expression level of *EhDHS1, EhDHS2, EheIF5A1, EheIF5A2*, and *EhRNA pol II* was expressed in arbitrary units. **(B)** Growth kinetics of control (pSAP2G) and *EhDHS2* gene silenced (DHS2gs) transformants. Approximately 6,000 amoebae in the logarithmic growth phase were inoculated into 6 mL fresh culture medium and amoebae were then counted every 24 h. Data shown are the means ± standard deviations of five biological replicates. **(C)** Immunoblot analysis of control (pSAP2G) and EhDHS2 gene silenced (DHS2gs) transformants using hypusine antibody. Total cell lysate was electrophoresed on a 15% SDS-PAGE gel and subjected to an immunoblot assay with hypusine antibody and CS1 (loading control) antiserum. The intensity of the bands corresponding to hypusinated EheIF5A and CS1 was measured by densitometric quantification, analyzed by Image J software, and is shown in arbitrary units in the right graph.

### Intracellular distribution of EhDHS1, EhDHS2, EheIF5A1, and EheIF5A2 and hypusine modification of EheIF5A proteins

To examine intracellular distribution of EhDHS1, EhDHS2, EheIF5A1, and EheIF5A2 in trophozoites, we established transformant lines expressing EhDHS1 and EhDHS2 with the hemagglutinin (HA) tag at the amino terminus (HA-EhDHS1, HA-EhDHS2, HA-EheIF5A1, and HA-EheIF5A2). The expression of these proteins in trophozoites was confirmed by immunoblot analysis with anti-HA antibody (S2 Fig). We first examined the distribution of EhDHS1, EhDHS2, EheIF5A1, and EheIF5A2 in trophozoites by immunoblotting using lysates produced by a Dounce glass homogenizer, and centrifugation at 5,000 *g* and subsequently at 100,000 *g*. We found that EhDHS1, EhDHS2, EheIF5A1, and EheIF5A2 were mainly found in the soluble fraction of centrifugation at 100,000 × g, together with a cytosolic soluble protein CS1 [28] (Fig 6A). We also found that EhDHS1 and EheIF5A1 (also in a lesser amount, EhDHS2 and EheIF5A2) were also associated with the pellet fraction after centrifugation at 100,000 g, together with a vesicular membrane protein CPBF1 [29] (Fig 6A). These data indicate that EhDHS1, EhDHS2, EheIF5A1, and EheIF5A2 were mainly present in the cytosol, but they are also partially localized to the some organelle(s)/compartments. Note that HA-EheIF5A1 is present as multiple bands, suggesting PTMs and those multiple forms are present in different proportions in different fractions

**Fig 6.**
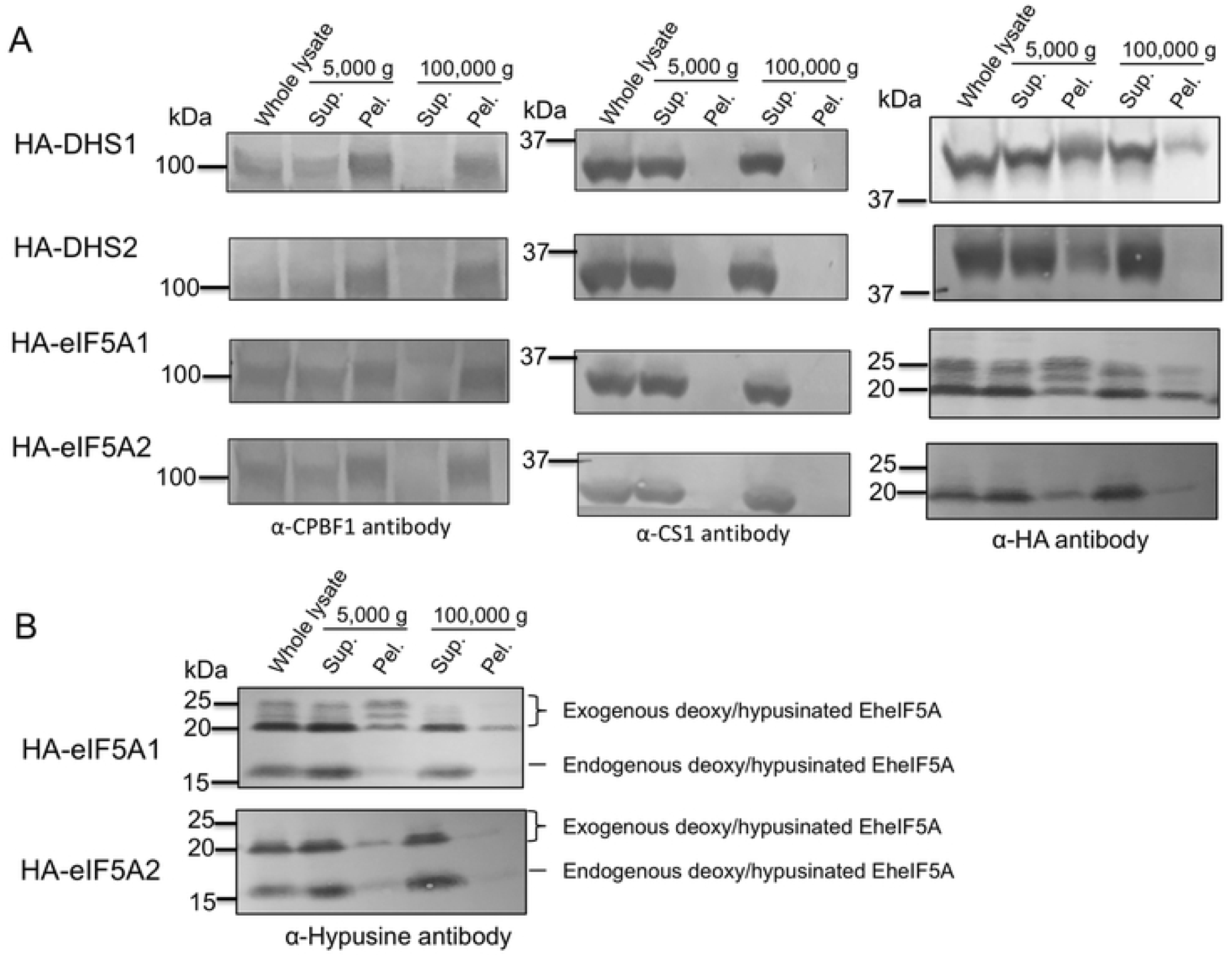
Cellular distribution of EhDHS and EheIF5A isoforms and post translational modification of EheIF5A1/2. **(A)** Cellular fractionation of EhDHS1, EhDHS2, EheIF5A1, and EheIF5A2. Trophozoites of the transformants expressing HA-tagged EhDHS1, EhDHS2, EheIF5A1, and EheIF5A2 were fractionated as described in Materials in Methods, and subjected to immunoblot analysis using anti-HA monoclonal antibody, anti-hypusine, anti-CPBF1, and anti-CS1 polyclonal antisera. CPBF1 and CS1 served as control of organelle and cytosolic proteins, respectively. **(B)** PTMs of HA tagged EheIF5A. Approximately 30 μg of total lysates from the transformants expressing HA-tagged EheIF5A1 and EheIF5A2 were subjected to SDS-PAGE under reducing conditions and immunoblot analysis using anti-hypusine antibody.

We further examined potential PTM of EheIF5A1 and EheIF5A2 using HA-EheIF5A1 and HA-EheIF5A2 overexpressing strains. Using anti-hypusine antibody, we detected four bands in HA-EheIF5A1-expressing trophozoites; the top three 20-25 kDa bands correspond to the exogenously expressed HA-EheIF5A1 and the bottom 17 kDa band corresponds to the endogenous EheIF5A (Fig 6B). In HA-EheIF5A2-expressing transformants, we detected two bands corresponding to one each of exogenous and endogenous EheIF5A (Fig 7A). However, we detected only a single band corresponding to the endogenous EheIF5A using hypusine antibody (Fig 6B). These data indicate that both EheIF5A1 and EheIF5A2 undergoes hypusination, but only EheIF5A1 is also subjected to PTMs other than hypusination. It was previously reported in other organisms that beside hypusination eIF5As are post-translationally modified by acetylation [21], sulfation [22], phosphorylation [30, 31] and glycosylation [31]. However, the nature of modifications in HA-EheIF5A1 expressing cells remains unknown. Our attempt to characterize PTMs on eIF5A1 by trypsinization and mass spectrometric analysis failed with repeated trials.

**Fig 7.**
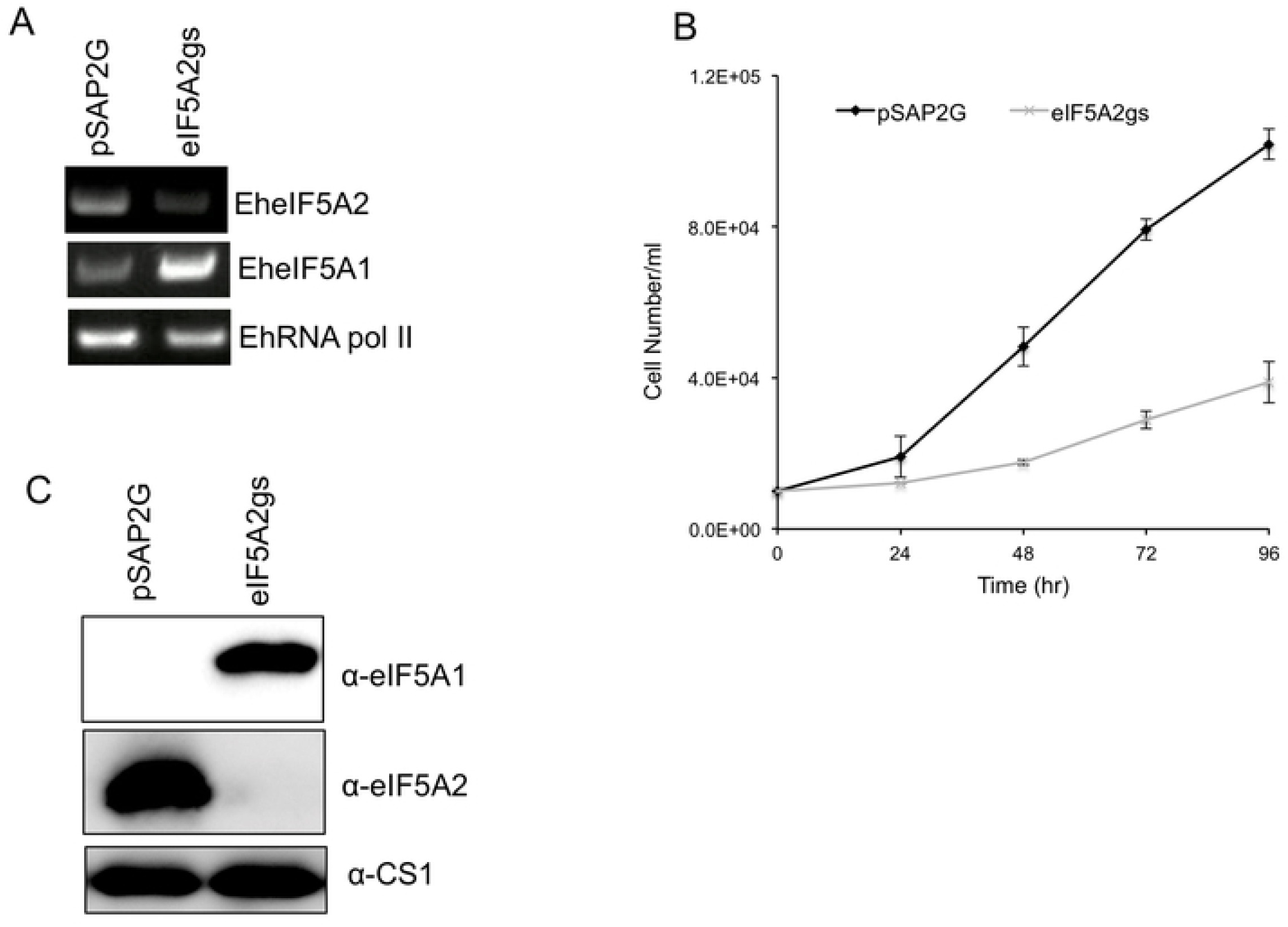
Silencing of *EheIF5A2* caused growth retardation and up regulation of EheIF5A expression. **(A)** Evaluation of gene expression by semi-quantitative RT-PCR analysis of *EheIF5A2* gene silenced transformant. The steady-state levels of transcripts of *EheIF5A1, EheIF5A2*, and *EhRNA pol II* genes were measured in trophozoites of G3 strain transfected with either empty vector (psAP2G) or the *EheIF5A2* gene silencing plasmid (psAP2G-eIF5A2). cDNA from the generated cell lines (psAP2G and EheIF5A2gs strains) was subjected to 30 cycles of PCR using specific primers for the *eIF5A1, eI5FA2* and *RNA pol II* genes. RNA polymerase II served as a control. PCR analysis of samples without reverse transcription was used to exclude the possibility of genomic DNA contamination. **(B)** Growth kinetics of *EheIF5A2* gene silenced (EheIF5A2gs) and control (pSAP2G) transformants. Approximately 6,000 amoebae in the logarithmic growth phase were inoculated into 6 mL fresh culture medium and amoebae were then counted every 24 h. Data shown are the means ± standard deviations of five biological replicates. **(C)** An immunoblot analysis of EheIF5A2gs and control (pSAP2G) transformants. Total cell lysates were electrophoresed on a 15% SDS-PAGE gel and subjected to an immunoblot assay with antibodies raised against EheIF5A1 and EheIF5A2.

Although EheIF5A hypusination in amebic trophozoites was verified by immunoblot analysis using anti-hypusine antibody, DOHH, which catalyzes the final step of the formation of the mature hypusinated eIF5A in other organism (Fig 1), seems to be missing in the *E. histolytica* genome (https://amoebadb.org/amoeba/). To verify if anti-hypusine polyclonal antibody used in this study recognizes hypusinated and/or deoxyhypusinated forms of eIF5A, we performed immunoblot analysis of recombinant eIF5A1 and eIF5A2 incubated in vitro with a combination of recombinant EhDHS1 and EhDHS2 (S3 Fig). We found that deoxyhypusinated form of eIF5A1 (lane 4) and eIF5A2 (lane 5) were detected using anti-hypusine antibody when they had been co-incubated with the recombinant DHSs and spermidine (S3 Fig). Thus, the nature of protein modification, that is, whether eIF5A is only deoxyhypusinated or fully hypusinated in amebic trophozoites, still remains elusive.

### Only eIF5A2 protein is expressed in trophozoites

In order to examine the levels of expression of each isoform of eIF5As in *E. histolytica* trophozoites, polyclonal antisera were raised against purified recombinant EheIF5A1 and EheIF5A2. The cell lysates of *E. histolytica* transformant cell lines expressing EheIF5A1 and EheIF5A2 with the HA tag fused at the amino terminus (HA-EheIF5A1 and HA-EheIF5A2) were analyzed by immunoblotting with antisera raised against EheIF5A1 and EheIF5A2. We confirmed specificities of these antisera using corresponding cell lines expressing EheIF5A1 and EheIF5A2 proteins (S4 Fig). To our surprise, only EheIF5A2, but not EheIF5A1, was detected in wild type trophozoites as well as all the transformants (S4 Fig). Our previous transcriptome analysis using DNA microarray [32] also suggests that *EheIF5A2* gene is abundantly transcribed as steady state mRNA while the mRNA levels of *EheIF5A1* gene is negligible.

### *EheIF5A2* gene silencing causes up regulation of *EheIF5A1* gene expression and also inhibits protein translation

In order to better understand the role of EheIF5A2, we utilized gene silencing to repress the *EheIF5A2* gene in *E. histolytica* G3 strain. *EheIF5A2* transcript was undetectable in EheIF5A2gs line (Fig 7A). Interestingly, we found that the steady-state level of *EheIF5A1* transcript was increased upon *EheIF5A2* gene silencing (Fig 7A). We further examined eIF5A1 and eIF5A2 expression at the protein level in EheIF5A2gs and control strains. Immunoblot analysis using anti-rEheIF5A1 and anti-rEheIF5A2 antibodies showed that eIF5A2 protein was undetected by EheIF5A2 antibody while eIF5A1 protein was detected in EheIF5A2gs strain, suggesting compensatory expression of EheIF5As for survival (Fig 7C). However, the compensatory upregulation of *eIF5A1* gene expression may not be sufficient to completely rescue the function of eIF5A2 because EheIF5A2gs strain still displayed growth defect (Fig 7B).

Since *EheIF5A2* gene silencing caused growth defect, we next examined whether it also affects protein synthesis per se. We utilized SUrface SEnsing of Translation (SUnSET) technique to monitor protein synthesis in *E. histolytica* [33]. To first validate that SUnSET is applicable to *E. histolytica*, amebic trophozoites were incubated with puromycin before or after incubation with cycloheximide, an inhibitor of de novo protein synthesis. Immunoblotting using anti-puromycin antibody revealed that puromycin was readily incorporated into proteins in *E. histolytica* (Fig 8A). The incorporation was dependent on de novo protein synthesis since no background staining was detected in the lysates from cyclohexamide treated cells (Fig 8A). We compared the protein synthesis efficiency in EheIF5A2gs and control strains. We found that in EheIF5A2gs strain, puromycin incorporation was almost completely abolished as compared to control (Fig 8B), indicating ceased protein synthesis caused by *EheIF5A2* gene silencing.

**Fig 8.**
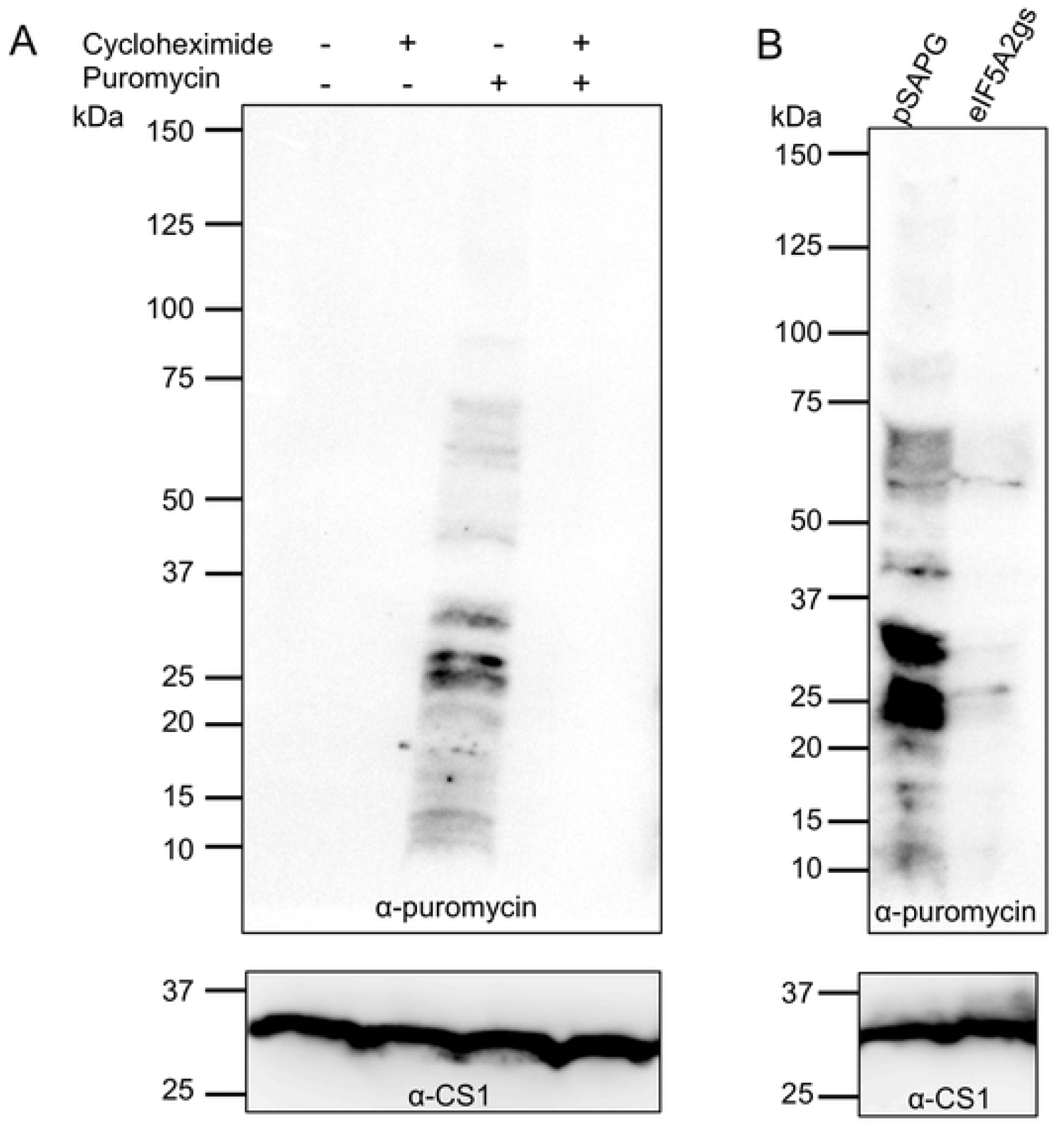
SUnSET analysis of active protein synthesis. **(A)** SUnSET analysis of wild type strain. Trophozoites of *E. histolytica* clonal strain HM-1:IMSS cl 6 were incubated in normal growth medium with 100 μg mL^-1^ cycloheximide, 10 μg mL^-1^ puromycin, or both. Cell lysates were subjected to SDS-PAGE and immunoblot using antibody specific for puromycin. Note that trophozoites incorporated puromycin into proteins (P). The incorporation was blocked by cycloheximide treatment (C + P). These data validated the SUnSET analysis for assessing protein synthesis in *E. histolytica*. **(B)** SUnSET analysis of EheIF5A2gs strain. Trophozoites from EheIF5A2gs and control (pSAP2G) strains were subjected to SUnSET analysis as in (A). Equal protein loads were confirmed with CS1 antibody (lower panels). Representative data for at least 3 separate trials are shown.

### eIF5A1 plays an important role in stage conversion

To get insights into the observations that only one *eIF5A* gene of two is expressed (eIF5A2) in trophozoites and to better understand the role of the other gene (*eIF5A1*). It was previously shown that eIF5A plays an important role in mouse embryogenesis and cell differentiation [34]. *Entamoeba* also undergoes differentiation between the proliferative trophozoite and the dormant cyst (encystation and excystation). We examined the transcriptomic profiles of *E. invadens*, a parasite for reptiles, which mimics biology and pathophysiology of *E. histolytica*, and easily undergoes encystation and excystation in vitro [35]. We first checked *E. invadens* genome database (https://amoebadb.org/amoeba/) and identified three eIF5A orthologs (EIN_344160, EIN_129200, and EIN_296010). We named them as EieIF5A1 (EIN_344160), EieIF5A2 (EIN_129200), and EieIF5A3 (EIN_296010) in an descending order of percentage amino acid identity with *E. histolytica* eIF5A1. EieIF5A1 showed 50% amino acid identity to EheIF5A1 and EieIF5A2 exhibit 73% amino acid identity to EheIF5A2, while EieIF5A3 shows only 23 and 15% amino acid identity to *E. histolytica* eIF5A1 and eIF5A2, respectively (S3 Table). A multiple alignment of three EieIF5A isotypes with two EheIF5A isoforms, by the ClustalW program (http://clustalw.ddbj.nig.ac.jp/top-e.html), shows that the two glycine residues adjacent to lysine (red) are also conserved in EieIF5A (Table S3). As mentioned above, this Gly-X-Y-Gly motif is reported to be critical for β turn structure and the proper orientation of the deoxyhypusine/hypusine side chain [20]. Lysine residues at amino acid positions 52 (EieIF5A1), 50 (EieIF5A2), and 66 (EieIF5A1) for hypusination are conserved in all three EieIF5A (Table S3). To determine whether eIF5As are involved in differentiation process of *E. invadens*, we carried out in vitro encystation of *E. invadens* using the methods described previously [36]. Approximately 90% of the trophozoites differentiated into the sarkosyl-resistant cysts within 120 hr (S5A Fig). Under the conditions, EieIF5A2 was abundantly expressed in *E. invadens* trophozoites similar to *E. histolytica*, while neither *EieIF5A1* nor *EieIF5A3* expression was detected in trophozoites (S5B Fig). In the course of encystation, the expression levels of *EieIF5A1-3* remained unchanged (S5B Fig), suggesting these genes are not involved in encystation of *Entamoeba*.

Next, we investigated whether eIF5As are involved in excystation. We found that approximately 45% and 88 % of the cysts transformed into trophozoites at 8 and 24 hr, respectively (Fig 9A). In the course of excystation, the steady-state level of *EieIF5A1* transcript was increased and peaked at 8 hr, while that of *eIF5A2* transcript was transiently abolished at 8 hr, but increased to the level similar to that in the cyst (Fig 9B), suggesting that eIF5A1 plays a excystation-specific role in *Entamoeba*. The transcript level of *EieIF5A3* gene, which is not expressed in trophozoites, remained unchanged throughout the excystation process.

**Fig 9.**
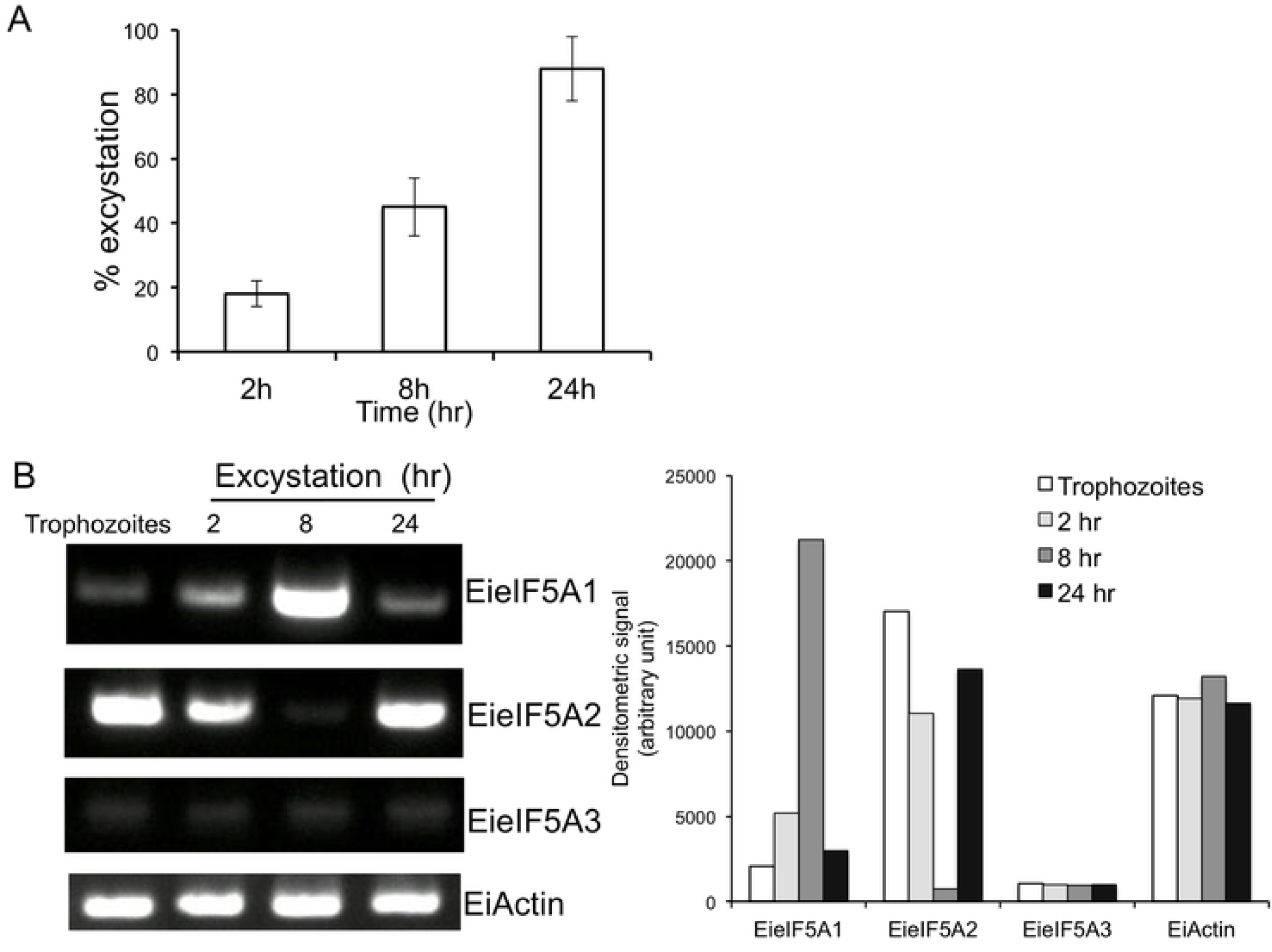
Steady state transcript levels of EieIF5A isoforms during excystation. **(A)** Kinetics of excystation. Cysts were incubated at the indicated times, cysts and trophozoites were harvested and the excystation percentage was determined. The excystation efficiency was determined by counting trophozoites and cysts in a haemocytometer and experiments were performed in duplicate. Data shown are the means ± standard deviations of two independent experiments. **(B)** Semi-quantitative RT-PCR analysis of *EieIF5A1, EieI5FA2, EieIF5A1* and *EiActin* genes during excystation. cDNA from 4 time points during encystation was subjected to 30 cycles of PCR using specific primers for *eIF5A1, eI5FA2, eIF5A3*, and *actin* genes. *Actin* gene served as a control. PCR from samples without reverse transcription served as controls to exclude the possibility of genomic DNA contamination. The densitometric quantification of the bands, shown in the right graph, was performed by Image J software, and the expression level of *EieIF5A1, EieIF5A2*, and *EieIF5A3 and EiActin* was expressed in arbitrary units.

## Discussion

### The first demonstration of hypusination of eIF5A and role of eIF5A on differentiation in *E. histolytica*

In this study, we have demonstrated that eIF5A and its PTMs are essential for both proliferation and differentiation in *Entamoeba*. Stage conversion is a key biological process needed for transmission of the eukaryotic pathogens [35-37]. We have shown that *Entamoeba* eIF5A is involved in excystation, conversion from the cyst to the trophozoite, but not encystation. *E. invadens*, used as an in vitro differentiation model, possesses three isoforms of eIF5A (S3 Table). Of two *E. invadens* isoforms orthologous to *E. histolytica* isoforms (EieIF5A1 and EieIF5A2), *EieIF5A1* gene was strongly upregulated only at 8 hours post excystation induction, strictly in a time-dependent fashion, when *EieIF5A2* expression was conversely repressed. In contrast, *EieIF5A2* gene was constitutively expressed in trophozoites, encysting trophozoites, and cysts. These data intriguingly indicate a specific role of EieIF5A1 in this very narrow time window of excystation. It needs to be elucidated in future how eIF5A1 and eIF5A2 are coordinately involved in excystation: whether eIF5A1 is involved in protein synthesis per se or other cellular functions, and in what cellular compartments. In humans, two eIF5A isoforms are known to exert specialized activities in certain tissues and cancer cells in addition to basic overlapping functions [38]. It was suggested that the divergent C-terminal domain of eIF5A isotypes may be involved in an interaction with alternate binding partners and responsible for homo-oligomerization as noted for human eIF5A1 [39] and archaeal counterpart [40].

### Catalytic and regulatory roles of two EhDHS isoforms

eIF5A is the only protein which undergoes hypusination in nature. No pathway is known for the synthesis of this modified amino acid. We have identified proteins responsible for deoxyhypusine modification of eIF5A, but failed to identify proteins responsible for the final enzyme to complete eIF5A hypusination. Whether DOHH is present in *E. histolytica* and, if so, its identity remain elusive. We have shown that two isoforms of EhDHS play catalytic and regulatory roles in eIF5A modifications. Most of eukaryotes possess either only a single DHS or two closely related active DHS. There are a few exceptions known other than *E. histolytica*: *Trypanosoma brucei* [16] and *Leishmania* [17], both of which, together with *E. histolytica*, possess two DHS, one catalytically active and one inactive proteins [16]. In *T. brucei*, it was reported that two DHS isotypes, two each of catalytically active and inactive isotypes formed a heterotetrameric complex, and DHS activity increased 3000-fold by heterotetramer formation [16]. It was also shown in *T. brucei* that loss of one of the isotypes led to instability of the complex, suggesting both DHS isotypes were necessary. Furthermore, both *EhDHS1* and *2* genes were essential for parasite growth [16]. In contrast to what was reported in *T. brucei*, we found that gene silencing of *EhDHS2* (inactive isoform) did not cause loss of either EhDHS1 protein or hypusinated eIF5A (Fig 5C). It is conceivable that in *EhDHS2* gene silenced strain, the decreased activity of EhDHS1 or instability of the EhDHS1/2 complex is partially compensated by transcriptional upregulation of *EhDHS1* and *eIF5A1*genes (Fig 5A).

### Possible posttranslational modifications of *E. histolytica* eIF5A other than hypusination

Hypusination was shown to play a critical role in the regulation of eIF5A [6]. In addition to hypusination, other modifications, such as phosphorylation [30, 31], glycosylation [31], acetylation [21, 41], transglutaminylation [42], and sulfation [22] of eIF5A have been described, but very little is known about their roles. In *E. histolytica*, both eIF5A1 and eIF5A2 are hypusinated, and only eIF5A1 undergoes other PTMs. It has been recently reported that human eIF5A rapidly translocates to the Golgi apparatus and is further modified by tyrosine-sulfation in the *trans*-Golgi prior to secretion [22]. Secreted tyrosine sulfated-eIF5A mediates oxidative stress-induced apoptosis by acting as a pro-apoptotic ligand [22]. It has been also reported in *T. vaginalis* that eIF5A is phosphorylated at serine 2 in a stretch of the consensus sequence present at the amino terminus (MSSAEEEV) [42]. A similar sequence (MSSNGSDN) is also present at the amino terminus of EheIF5A2, but not in EheIFA1, suggesting that EheIF5A2 may be phosphorylated (Fig 2B). It has been reported that in maize, a serine 2 the amino terminus (MSDSEEHH) of eIF5A is phosphorylated by the catalytic α-subunit of the casein kinase 2 (CK2) [30]. This phosphorylation is also known to play a role in the regulation of the nucleo-cytoplasmic shuttling of eIF-5A in plant cells [44]. However, we failed to detect phosphorylation of EheIF5A2 using phospho serine antibody by immunoblot analysis (data not shown).

### Localization of eIF5A may be attributable to posttranslational modifications

We found by cellular fractionation that EhDHS1, EhDHS2, EheIF5A1, and EheIF5A2 are associated with the 5,000 g pellet and 100,000 g pellet fractions, which contain the nucleus, the plasma membrane, large vacuoles (the former fraction), mitosomes, the endoplamsic reticulum, and lysosomes (the latter), although these four proteins were mostly associated with the soluble cytosolic fraction (Fig 6). Intriguingly, three hypusinated bands corresponding to EheIF5A1 were mainly detected in the 5000 g membrane fraction using anti-hypusine antibody, and the band patterns of hypusinated EheIF5A1 differed in 5,000 g pellet and other fractions (note the middle band is only detected in 5,000 g pellet), suggesting that the those heterogeneously modified eIF5A1 proteins are associated with the nucleus, the plasma membrane, and/or heavy membranes. Such PTMs on EheIF5A1 seem to modulate its functions by targeting EheIF5A1 to different cellular compartments to impair the interaction with its partners as demonstrated for human eIF5A [45]. The subcellular localization of eIF5A and its nuclear-cytoplasmic shuttling in other organisms remain controversial [46]. It has been reported that eIF5A is localized to the ER in multiple mammalian cell lines [47], while other studies have shown that eIF5A shuttles between the nucleus and the cytoplasm, and occasionally co-localizes with the nuclear pore complex [48]. It was also demonstrated that eIF5A enters the nucleus through passive diffusion and is exported from the nucleus to the cytoplasm in a CRM1(chromosomal maintenance 1, also known as exportin1)-dependent manner [49]. It was also shown that exportin 4, an importin β family receptor, is responsible for the nuclear export of eIF5A [50,51]. Genome survey of *E. histolytica* suggest that exportin 1 (EHI_107080) and exportin T (EHI_029040) are encoded by the genome. It has been recently reported that exogenous eIF5A lacking hypusination tends to localize to the nucleus, compared to fully hypusinated endogenous eIF5A that mainly localizes to the cytoplasm, but can be translocated to the cytoplasm when fully hypusinated [52], suggesting that hypusination promotes a nucleus-to-cytoplasm transport. Nevertheless, our immunofluorescence imaging failed to detect nuclear localized eIF5A (S6 Fig).

### Multiplicity and essentiality of eIF5A

We showed that eIF5A2, but not eIF5A1, is constitutively expressed in trophozoites, although both the isoforms harbor the hypusine modification (Fig 7C). Two or more *eIF5A* genes are present in various eukaryotes including fungi, plants, vertebrates, and mammals [53, 54]. In fish, amphibians, and birds, the two *eIF5A* genes seem to be coexpressed [55]. In humans, one eIF5A isoform is abundantly expressed in most cells, essential for cell proliferation, while the other isoform is undetectable or only weakly expressed in most cells and tissues, but highly expressed in various cancerous cells, and thus suggested to be associated with cancers [56, 57].

Gene disruption and mutation studies in yeast and higher eukaryotes indicate the essentiality of eIF5A and its deoxyhypusine/hypusine modifications [58]. In *Saccharomyces cerevisiae*, simultaneous disruption of two *eIF5A* genes or inactivation of the single *DHS* gene led to growth arrest, and substitution of one of two eIF5As with eIF5A K50R mutant also caused growth arrest [59, 60]. Interestingly, *EheIF5A2* gene silencing resulted in the compensatory up regulation of *EheIF5A1* gene expression (Fig 7A and C), suggesting role of eIF5A1 and eIF5A2 is interchangeable to some extent. Despite the observed *eIF5A1* upregulation, *EheIF5A2* gene silenced strain showed defect in protein synthesis, suggesting eIF5A2 also plays unique role in protein synthesis and proliferation (Fig 8B). The mechanisms by which *EheIF5A2* gene silencing causes upregulation of *EheIF5A1* gene expression remains elusive. Genetic compensation has been documented in a number of model organisms [61]. It has been proposed that in higher eukaryotes, mRNA surveillance pathways, small non-coding RNAs (ncRNAs), upstream open reading frames (uORFs), RNA-binding proteins (RBPs), and micro-RNAs (miRNAs) can potentially be involved in the compensatory response [61].

In conclusion, we have demonstrated that eIF5A and its PTMs are essential for proliferation and differentiation of *Entamoeba*. Furthermore, we have shown that two isoforms of EhDHS play catalytic and regulatory roles in eIF5A modifications. Our results demonstrated for the first time a clear reason for the significance for spermidine, the substrate for hypusination for *Entamoeba* biology. Our present data should provide a rationale for the PTM of translational machinery and the biosynthetic pathway of polyamines are a good target for the development of new drugs against *Entamoeba*. The specific roles of the unique modifications on EheIF5A1 and identification of DOHH and the enzymes involved in polyamine biosynthesis need to be elucidated in future.

## Materials and methods

### Chemicals and Reagents

NAD^+^, spermidine, and 1,3-diaminopropane were purchased from Sigma–Aldrich (Tokyo, Japan). Ni^2+^-NTA agarose was purchased from Novagen (Darmstadt, Germany). Lipofectamine and geneticin (G418) were purchased from Invitrogen (Carlsbad, CA, USA). All other chemicals of analytical grade were purchased from Wako (Tokyo, Japan) unless otherwise stated.

### Microorganisms and cultivation

Trophozoites of the *E. histolytica* clonal strain HM-1:IMSS cl6 and G3 strain [62] were maintained axenically in Diamond’s BI-S-33 medium at 35.5°C as described previously [63]. Trophozoites were harvested in the late-logarithmic growth phase for 2–3 days after inoculation of one-thirtieth to one-twelfth of the total culture volume. After the cultures were chilled on ice for 5 min, trophozoites were collected by centrifugation at 500 × g for 10 min at 4°C and washed twice with ice-cold PBS, pH 7.4. *Escherichia coli* BL21 (DE3) strain was purchased from Invitrogen.

### Alignment of DHS and eIF5A protein sequences

Amino acid sequences of DHS protein from *E. histolytica* and other organisms were aligned to examine sequences surrounding the key lysine reside. Similarly, amino acid sequences of eIF5A protein from *E. histolytica* and human were aligned to examine the key lysine residue that is post translationally modified by hypusination and acetylation. Multiple sequence alignments were generated using CLUSTAL W program (http://clustalw.ddbj.nig.ac.jp/) [64].

### Construction of plasmids for the production of recombinant *E. histolytica* DHS1, DHS2, eIF5A1, and eIF5A2

Standard techniques were used for cloning and plasmid construction, essentially as previously described [65]. A DNA fragment corresponding to cDNA encoding *Eh*DHS1, *Eh*DHS2, *Eh*eIF5A1, and *Eh*eIF5A2 was amplified by PCR from *E. histolytica* cDNA using the oligonucleotide primers listed in Table S1 to produce a fusion protein containing a histidine-tag (provided by the vector) at the amino terminus. PCR was performed with platinum *pfx* DNA polymerase (Invitrogen) and the following parameters: an initial incubation at 94 °C for 2 min; followed by the 30 cycles of denaturation at 94 °C for 15 s; annealing at 50, 45, or 55 °C for 30 s; and elongation at 68 °C for 2 min; and a final extension at 68 °C for 10 min. The PCR fragment was digested with *Bam*HI and *Sal*I, electrophoresed, purified with Gene clean kit II (BIO 101, Vista, CA, USA), and ligated into *Bam*HI and *Sal*I digested pCOLD-1 (Novagen) in the same orientation as the T7 promoter to produce pCOLD1-EhDHS1, EhDHS2, EheIF5A1, and EheIF5A2. The nucleotide sequence of the cloned EhDHS1, DHS2, eIF5A1, and eIF5A2 was verified by sequencing to be identical to the putative protein coding region in the genome database (http://amoebadb.org/amoeba/).

### Construction of a plasmid for coexpression of EhDHS1 and EhDHS2

To construct a plasmid for EhDHS1 and EhDHS2 coexpression, *EhDHS1* and *EhDHS2* genes were amplified using PCR. The upstream and downstream oligonucleotide primers of *EhDHS1* gene, listed in Table S1, contained a *Bam*HI restriction site and a *Hin*dIII restriction site. The PCR fragment was subsequently cloned into the *Bam*HI-*Hin*dIII restriction sites of pETDuet (Novagen) to generate the recombinant plasmid pETDuet-DHS1. The His-Tag sequence in pETDuet was fused with *EhDHS1* gene at the N-terminus. The upstream and downstream oligonucleotide primers of *EhDHS2* gene, listed in Table S1, contained *Aat*II and *Xho*I restriction sites. The stop codon of *EhDHS2* gene was removed so that *EhDHS2* gene is fused with the region encoding the C-terminal S-Tag in pETDuet. The PCR fragment was subsequently cloned into the *Aat*II-*Xho*I restriction sites of pETDuet-*Eh*DHS1 to generate pETDuet-*Eh*DHS1/2.

### Site directed mutagenesis of EhDHS1 and EhDHS2

One-step site directed mutagenesis was conducted on *EhDHS1* and *EhDHS2* gene by QuikChange Site-Directed Mutagenesis Kit (Agilent Technologies). Overlapping primers for site-directed mutagenesis were designed to replace phenylalanine at amino acid 302 of the pCOLD-EhDHS2 plasmid with lysine. Similarly, lysine at amino acid 295 of the pCOLD-EhDHS1 plasmid was replaced with phenylalanine. The primer sets used are listed in Table S1. PCR reaction was carried out using 100 ng/μl each of the primers, 200 mM dNTPs, 2U of *PfuTurbo* DNA polymerase (Vendor, City, Country) and pCOLD-EhDHS1 or pCOLD-EhDHS2 as a template. The PCR parameters were: 95°C for 30 sec; 16 cycles of 95°C for 30sec, 55°C for 1 min and 68°C for 10 min. The PCR-amplification products were evaluated by agarose gel electrophoresis. The PCR product were digested with 1 μl of the *Dpn* I restriction enzyme (10 U/μl) at 37°C for 1 hour to digest the parental (i.e., the non-mutated) supercoiled dsDNA. Approximately 1 μl of the *Dpn* I-treated DNA from each sample reaction was transferred to a tube containing *E. coli* XL1-Blue cells. The tube was incubated for 45 seconds at 42°C, and then placed on ice for 2 minutes. The cells were spread onto an LB plate containing 100 μg/mL ampicillin. Bacterial clones containing the plasmid that encodes mutated EhDHS1 or EhDHS2 (DHS1_K295F, DHS2_F302K, respectively) that possesses the intended amino acid mutation (phenylalanine at amino acid 295 in pCOLD-EhDHS1 and lysine at amino acid 302 in pCOLD-EhDHS2) were selected by restriction enzyme digestion and DNA sequencing of the plasmids.

### Bacterial expression and purification of recombinant EhDHS1, EhDHS2, EhDHS1_K295F, EhDHS2_F302K, EheIF5A1, and EheIF5A2

The above mentioned plasmids were introduced into *E. coli* BL21 (DE3) cells by heat shock at 42°C for 1 min. *E. coli* BL21 (DE3) strain harboring pCOLD1-EhDHS1, EhDHS2, EheIF5A1, EheIF5A2, EhDHS1_K295F, EhDHS2_F302K, or pETDuet-EhDHS1/2 was grown at 37 °C in 100 ml of Luria Bertani medium in the presence of 50 μg/ml ampicillin. The overnight culture was used to inoculate 500 ml of fresh medium, and the culture was further continued at 37°C with shaking at 180 rpm. When A_600_ reached 0.6, 1mM of isopropyl ß-d-thio galactopyranoside was added, and cultivation was continued for another 24 h at 15°C except pETDuet-EhDHS1/2 which cultivated for another 6 h at 30°C. *E. coli* cells from the induced culture were harvested by centrifugation at 4,000 × g for 20 min at 4°C. The cell pellet was washed with PBS, pH 7.4, re-suspended in 20 ml of the lysis buffer (50 mM Tris–HCl, pH 8.0, 300 mM NaCl, and 10 mM imidazole) containing 0.1% Triton X100 (v/v), 100 μg/ml lysozyme, and 1 mM phenylmethyl sulfonyl fluoride, and incubated at room temperature for 30 min, sonicated on ice and centrifuged at 25,000 × g for 15 min at 4 °C. The supernatant was mixed with 1.2 ml of 50% Ni^2+^-NTA His-bind slurry, incubated for 1 h at 4°C with mild shaking. The resin in a column was washed three times with buffer A [50 mM Tris-HCl, pH 8.0, 300 mM NaCl, and 0.1% Triton X-100, v/v] containing 10-50 mM of imidazole. Bound proteins were eluted with buffer A containing 100-300 mM imidazole to obtain recombinant EhDHS1, EhDHS2, a combination of EhDHS1 and 2, EhDHS1_K295F, EhDHS2_F302K, EheIF5A1, and EheIF5A2. After the integrity and the purity of recombinants protein were confirmed with 12% SDS-PAGE analysis, followed by Coomassie Brilliant Blue staining, they were extensively dialyzed twice against the 300-fold volume of 50 mM Tris-HCl, 150 mM NaCl, pH 8.0 containing 10% glycerol (v/v) and the Complete Mini protease inhibitor cocktail (Roche, Mannheim, Germany) for 18 h at 4 °C. The dialyzed proteins were stored at −80 °C with 30% glycerol in small aliquots until further use. Protein concentrations were spectrophotometrically determined by the Bradford method using bovine serum albumin as a standard as previously described [66].

### Enzyme assays

DHS enzymatic activity was determined by the method described previously [23], unless otherwise stated. The duration of the reaction, buffer pH, and the concentrations of the enzyme and substrates were optimized for the assay. Briefly, a reaction mixture of 50 μl containing 0.2 M glycine-NaOH buffer, pH 9.2, 1 mM dithiothreitol, 1 mM NAD^+^, 1mM spermidine, 5 μM eIF5A, and the 3μg of recombinant EhDHS1 or 2 was incubated at 37°C for 60 min and reaction was terminated by the addition of ice cold 10% trichloroacetic acid. The samples were then centrifuged at 10,000 × g for 5 min at 4°C and the clear supernatant was analyzed for 1,3 diaminopropane production using reverse phase high performance liquid chromatography as described previously [67].

### Generation of *E. histolytica* transformants overexpressing epitope-tagged EhDHS1, EhDHS2, EheIF5A1, EheIF5A2

The protein coding region of *EhDHS1/2* and *EheIF5A1/2* genes were amplified from cDNA by PCR using sense and antisense oligonucleotide primers that contain *Sma*I and *Xho*I sites at the 5’ end, listed in Table S1. The PCR-amplified DNA fragment was digested with *Sma*I and *Xho*I, and ligated into *Sma*I- and *Xho*I-double digested pEhExHA [68] to produce pEhExHA-DHS1, DHS2, eIF5A1, and eIF5A2. Wild-type trophozoites were transformed with pEhExHA-EhDHS1, pEhExHA-EhDHS2, pEhExHA-EheIF5A1, and pEhExHA-EheIF5A1 by liposome-mediated transfection as previously described [69]. Transformants were initially selected in the presence of 3 μg/ml of geneticin. The geneticin concentration was gradually increased to 6-10 μg/ml during next 2 weeks before the transformants were subjected to analyses.

### Production of *EhDHS1/2* and *EheIF5A1/A2* gene-silenced strains

In order to construct plasmids for small antisense RNA-mediated transcriptional gene silencing [62] of *EhDHS1/2* and *EheIF5A1/A2* genes, a fragment corresponding to a 420-bp long fragment corresponding to the amino-terminal portion of the entire protein was amplified by PCR from cDNA using sense and antisense oligonucleotide primers containing *Stu*I and *Sac*I restriction sites (restriction sites are shown in bold), as shown in Table S1. The PCR-amplified product was digested with *Stu*I and *Sac*I, and ligated into the *Stu*I- and *Sac*I-double digested psAP2-Gunma [70] to construct gene silencing plasmids psAP2G-DHS1, psAP2G-DHS2, psAP2G-eIF5A1 and psAP2G-eIF5A2. The trophozoites of G3 strain were transformed with either empty vector or silencing plasmid by liposome-mediated transfection as previously described [69]. Transformants were initially selected in the presence of 1 μg/ml geneticin, and the geneticin concentrations were gradually increased to 6-10 μg/ml during next two weeks prior to subjecting the transformants to analyses.

### Growth assay of *E. histolytica* trophozoites

Approximately 6×10^6^ exponentially growing trophozoites of *E. histolytica* G3 strain transformed with psAP2G-DHS2, psAP2G-eIF5A2, and psAP2G (control) were inoculated into 6 ml of fresh BI-S-33 medium containing 10 μg/mL geneticin, and the parasites were counted every 24 h on a haemocytometer.

### Surface sensing of translation (SUnSET)

SUnSET analysis was conducted to assess translational machinery in *E. histolytica* as previously described [33]. Approximately 2×10^6^ trophozoites were incubated with 10 μg/ml puromycin (Sigma-Aldrich) for 15 min before or after incubation with 100 μg/ml cycloheximide for 10 min at 37°C. After cells were collected by centrifugation, proteins were precipitated by incubating with 20% (v/v) TCA on ice for 10 min, followed by centrifugation at 2200 × g for 5 min and washing the pellet with 5% (v/v) TCA. The protein pellet was resuspended in 2× SDS running buffer and incubated in boiling water for 10 min. The lysate was frozen at −80°C until analyzed via immunoblot analysis (see below). Anti-puromycin mouse monoclonal antibodies (Sigma-Aldrich) were used at a 1:2500 dilution. As a loading control, blots were reacted with anti-cysteine synthase 1 (CS1) rabbit polyclonal antiserum [28] at a 1:1000 dilution.

### *E. invadens* culture, encystation, and excystation

Trophozoites of *E. invadens* IP-1 strain were cultured axenically in BI-S-33 medium at 26°C. To induce encystation, 2 week-old *E. invadens* cultures were passaged to 47% LG medium lacking glucose [13, 36] at approximately 6×10^5^ cells/ml. Amebae were collected at various time points, and the formation of cysts was assessed in triplicate by virtue of the resistance to 0.05% sarkosyl using 0.22% trypan blue to selectively stain dead cells. Cysts were also verified by cyst wall staining by incubating amebae with calcofluor white (fluorescent brightener; Sigma-Aldrich) at room temperature. For excystation, cysts formed as above were harvested at 72 hr and any remaining trophozoites were destroyed by hypotonic lysis by incubating overnight (∽16 hr) in distilled water at 4°C. Excystation was induced by incubating 8×10^5^ cysts in 6 ml of LG medium supplemented with 1 mg/ml bile, 40 mM sodium bicarbonate, 1% glucose, 2.8% vitamin mix, and 10% adult bovine serum for 48 hr [71]. Excystation was conducted in duplicates and the efficiency was determined by counting trophozoites on a haemocytometer.

### Cell fractionation and immunoblot analysis

Trophozoites of the ameba transformant expressing pEhExHA-DHS1, pEhExHA-DHS2, pEhExHA-eIF5A1, or pEhExHA-eIF5A1, were washed three times with PBS containing 2% glucose. After resuspension in homogenization buffer (50 mM Tris-HCl, pH 7.5, 250 mM sucrose, 50 mM NaCl and 0.5 mg/ml E-64 protease inhibitor), the cells were disrupted mechanically by a Dounce homogenizer on ice, centrifuged at 500 × g for 5 min, and the supernatant was collected to remove unbroken cells. The supernatant fraction was centrifuged at 5000 × g for 10 min to isolate pellet and supernatant fractions. The 5000 × g supernatant fraction was further centrifuged at 100,000 × g for 60 min to produce a 100,000 × g supernatant and pellet fractions. The pellets at each step were further washed twice with homogenization buffer and re-centrifuged at 100,000 × g for 10 min to minimize carryover.

Cell lysates and culture supernatants were separated on 12-15% (w/v) SDS-PAGE and subsequently electro transferred onto nitrocellulose membranes (Hybond-C Extra; Amersham Biosciences UK, Little Chalfont, Bucks, UK) as previously described [72]. Non-specific binding was blocked by incubating the membranes for 1.5 h at room temperature in 5% non-fat dried milk in TBST (50 mM Tris-HCl, pH 8.0, 150 mM NaCl and 0.05% Tween-20). The blots were reacted with anti-HA mouse monoclonal (clone 11MO, Covance, Princeton, NJ, USA), anti-hypusine rabbit polyclonal antibodies [27], anti-cysteine protease binding family protein 1 (CPBF1) [29], and anti-cysteine synthase 1 (CS1) rabbit polyclonal antisera [29] at a dilution of 1:500 to 1:1000. CPBF1 and CS1 served as controls for membrane and cytosolic fractions, respectively. The membranes were washed with TBST and further reacted either with alkaline phosphatase-conjugated anti-mouse or anti-rabbit IgG secondary antibody (1:2000) or with horse radish peroxidase-conjugated anti-mouse or anti-rabbit IgG antisera (1:20,000) (Invitrogen) at room temperature for 1 h. After washings with TBST, specific proteins were visualized with alkaline phosphatase conjugate substrate kit (Bio-Rad) and images were scanned with Image Scanner (Amersham Pharmacia Biotech, Piscataway, NJ, USA) or the fluorescent signal of each protein was measured with a chemiluminescence detection system (Millipore) using Scion Image software (Scion Corp., Frederick, MD).

## Acknowledgements

We acknowledge all members of Nozaki lab for invaluable input, especially Kumiko Nakada-Tsukui, Somlata, Koushik Das, and Herbert Santos for technical assistance and valuable discussions.

## Author Contributions

Conceived and designed the experiments: GJ and TN. Performed the experiments: GJ. Analyzed the data: GJ. Contributed reagents/materials/analysis tools: TN. Wrote the paper: GJ and TN.

## Supplementary information

**S1 Table**. List of primers used in this study.

**S2 Table**. Multiple alignment of amino acid sequences of four entries corresponding to *E. histolytica* eIF5A. Accession numbers of these sequences are as follows: EHI_177460 (XP_655916), EHI_151810 (XP_657397), EHI_151540 (XP_657374), and EHI_186480 (XP_651531). The conserved residues are marked by asterisks (*) while similar amino acids are shown either with periods (.) or colons (:). The conserved lysine residue, which was supposed to be hypusinated, is highlighted in red. Sequence alignment was performed using ClustalW.

**S3 Table. (A)** Percent amino acid identity among *E. histolytica* and *E. invadens* eIF5A isoforms by ClustalW multiple sequence alignment score. GenBank accession numbers: EheIF5A1 (XP_657374), EheIF5A2 (XP_651531), EieIF5A1 (XP_004255257), EieIF5A2 (XP_004258351), EieIF5A3 (XP_004260381). **(B)** Amino acid sequence alignment of *E. histolytica* and *E. invadens* eIF5A isoforms. The conserved residues are marked by asterisks (*) while similar amino acids are shown either with periods (.) or colons (:). The conserved lysine residue, which was supposed to be hypusinated, is highlighted in red. Sequence alignment was performed using ClustalW.

**S1 Fig. (A) Purification of recombinant EhDHS1, EhDHS2, EheIF5A1, and EheIF5A2**. Protein samples at each step of purification were subjected to 15% SDS-PAGE under reducing conditions, and then stained with Coomassie Brilliant Blue R250. **(B)** Purification of the recombinant EhDHS mutants.

**S2 Fig**. Immunoblot analysis of *E. histolytica* transformant strains expressing HA-tagged EhDHS1, EhDHS2, EheIF5A1 and EheI5A2. Approximately 40 μg of total lysates were electrophoresed on a SDS-PAGE gel under reducing conditions and subjected to immunoblot analysis using anti-HA and anti-hypusine antibodies.

**S3 Fig**. In vitro deoxyhypusination of EheIF5A by coexpressed recombinant EhDHS1 and 2, and verification of the specificity of anti-hypusine antibody. Recombinant EheIF5A1 or EheIF5A2 was incubated with 1 mM spermidine, 0.5 mM NAD^+^, 3μg of coexpressed recombinant EhDHS1 and 2 (N-terminal His-tag EhDHS1 and C-terminal S-tag EhDHS2), and the mixtures were subjected to SDS-PAGE and immunoblot analyses using anti-hypusine, His-tag, and S-tag antibodies.

**S4 Fig. Validation of expression of Endogenous and epitope tagged EheIF5A isoforms in *E. histolytica* trophozoites**. Approximately 40 mg of total lysate from the transformant expressing HA-EheIF5A1 and HA-EheIF5A2 and HA was electrophoresed on a SDS-PAGE gel under reducing conditions and subjected to immunoblot analysis using anti-EheIF5A1, EheIF5A2 and anti-HA antibodies.

**S5 Fig. The levels of steady state mRNA expression of *EheIF5A* isoforms during encystation. (A)** Kinetics of encystation. The percentages of the amoebae resistant to 0.05% sarkosyl during encystation. **(B)** The steady-state levels of transcripts of *EieIF5A1, EieI5FA2, EieIF5A1*, and *EiActin* genes measured by semi-quantitative RT-PCR during encystation. cDNA from different time points during encystation was subjected to 30 cycles of PCR using specific primers for the *EheIF5A1, EheI5FA2, EheIF5A3* and *Ehactin* genes. *Ehactin* gene served as a control. PCR analysis of samples without reverse transcription was also used to exclude the possibility of genomic DNA contamination.

**S6 Fig**. Representative immunofluorescence assay (IFA) micrographs of HA-eIF5A1 and HA-eIF5A2 expressed in *E. histolytica* trophozoites, double stained with anti-HA antibody (green) and anti-CS1 antiserum (red) respectively. EhCS1 (Cysteine synthase 1) served as a cytosolic control. Scale bar, 10 μm.

